# Pharmacological modulation of the cytosolic oscillator affects glioblastoma cell biology

**DOI:** 10.1101/2023.11.23.568279

**Authors:** Paula M Wagner, Mario E Guido

**Affiliations:** CIQUIBIC-CONICET, Facultad de Ciencias Químicas, Universidad Nacional de Córdoba, Córdoba, Argentina; Departamento de Química Biológica Ranwel Caputto, Facultad de Ciencias Químicas, Universidad Nacional de Córdoba, Córdoba, Argentina

**Author notes:** Corresponding Authors –.

**Keywords:** circadian rhythm, glioblastoma, chronotherapy, metabolic oscillator

## Abstract

The circadian system is a conserved time-keeping machinery that regulates a wide range of processes such as sleep/wake, feeding/fasting, and activity/rest cycles to coordinate behavior and physiology. Circadian disruption can be a contributing factor in the development of metabolic diseases, inflammatory disorders, and higher risk of cancer. Glioblastoma (GBM) is a highly aggressive grade 4 brain tumor that is resistant to conventional therapies and has a poor prognosis after diagnosis, with a median survival of only 12-15 months. GBM cells kept in culture were shown to contain a functional circadian oscillator. In seeking more efficient therapies with lower side effects, we evaluated the pharmacological modulation of the circadian clock by targeting the cytosolic kinases glycogen synthase kinase-3 (GSK-3) and casein kinase ε/δ (CK1ε/δ) with specific inhibitors (CHIR99022 and PF670462, respectively), the cryptochrome protein stabilizer (KL001), or circadian disruption after *Per*2 knockdown expression in GBM-derived cells. CHIR99022-treated cells had a significant effect on cell viability, clock protein expression, migration, and cell cycle distribution. Moreover, cultures exhibited higher levels of reactive oxygen species and alterations in lipid droplet content after GSK-3 inhibition as compared with control cells. The combined treatment of CHIR99022 with temozolomide was found to improve the effect on cell viability compared to temozolomide therapy alone. *Per*2 disruption affected both GBM migration and cell cycle progression. Overall, our results suggest that pharmacological modulation or molecular clock disruption severely affects glioblastoma cell biology.

## INTRODUCTION

The circadian system present in all vertebrates, including mammals, temporally regulates behavioral, physiological, and molecular rhythms with a period close to 24 hours, allowing organisms to anticipate the environmental cycles with implications for health and disease [1]. The system comprises the central clock located in the suprachiasmatic nucleus (SCN) of the hypothalamus and peripheral clocks present in different tissues throughout the body. The central clock receives light, the main synchronizer of the circadian system, through intrinsically photosensitive retinal ganglion cells and is thus able to coordinate physiology and behavior through outputs sent to the peripheral clocks. These circadian oscillators are present even in individual cells and constitute highly conserved molecular machinery in diverse species [2–4]. At the molecular level, the circadian machinery involves transcriptional– translational feedback loops (TTFLs) responsible for generating the ∼ 24-hour oscillations in gene expression. In the main loop, the positive elements of the circuit, BMAL1 and CLOCK, heterodimerize and recognize E-box sequences in the promoter of the period (*Per*1, *Per*2, *Per*3) and cytochrome (*Cry*1, *Cry*2) genes to activate their transcription. Once the negative regulators PER and CRY accumulate in the cytoplasm and heterodimerize, they translocate to the nucleus to repress their own transcription by interacting with the BMAL1: CLOCK complex [5–8]. Degradation of the negative regulators also plays a role in closing a 24-hour cycle and starting a new cycle. The secondary loop includes the nuclear receptors REV-ERB α/β and ROR α/β/γ, which recognize retinoic acid receptor-related orphan receptor response elements (RORE) in the promotor region of target genes. In turn, REV-ERB proteins repress *Bmal*1 and *Npas*2 transcription whereas ROR proteins can activate *Clock* and *Bma1*1 expression [9–12]. In light of recent evidence of PER2 protein oscillation in a CRY-deficient model, Putker and colleagues suggest that the TTFL is required for the robustness of behavioral and physiological rhythmicity but is dispensable for circadian timekeeping per se. This suggests the existence of an alternative coupled underlying timekeeping mechanism termed “cytoscillator” involving the casein kinase δ/ɛ (CK1δ/ɛ) and glycogen synthase kinase 3 (GSK-3) [13]. This post-translational mechanism is evolutionarily conserved across a range of model systems including isolated red blood cells (reviewed in [14, 15]), and can function independently of canonical clock proteins [16]. GSK-3 is a serine/threonine protein kinase with the two functional isoforms α and β. The latter isoform is a protein kinase of 47 kDa that regulates cellular processes such as proliferation, cell cycle control, migration, invasion, and apoptosis [16–18]. Evidence in the literature suggests that GSK-3 phosphorylates at least five clock proteins such as PER2, CRY2, CLOCK, BMAL1, and REVERBα [19, 20]. Previously, it was shown that the selective pharmacological inhibition of GSK-3 by CHIR99021, a synthetic aminopyrimidine derivative, accelerates the speed of the cellular timekeeping system in a wide range of organisms even in the absence of transcription [13, 17–20]. Moreover, CHIR99021 treatment resulted in reduced cell proliferation of epithelioid sarcoma [21] and tumor growth inhibition when combined with paclitaxel in non-small cell lung carcinoma [22]. Although circadian disruption could alter metabolic pathways that lead to pathologies, little is known about the pharmacological modulation of cytosolic kinases as a potential therapeutic strategy to treat brain cancer. Based on the World Health Organization (WHO) classification, GBMs are grade 4 brain tumors resistant to conventional therapies; they represent 45.2% of all malignant central nervous system (CNS) tumors and 80% of all primary malignant CNS tumors [23]. The standard of care treatment is the Stupp protocol, which combines radiotherapy followed by the administration of the DNA-alkylating chemotherapeutic temozolomide (TMZ) (reviewed in [24]). Though the TMZ regimen was approved by the US Food and Drug Administration (FDA), its chemotherapeutic efficiency is poor and most patients relapse [25]. GSK-3β is expressed in all tissues, with its highest expression in the developing brain [26]. Miyashita and collaborators reported higher expression levels and activity of GSK-3β in GBM compared with normal brain tissues [27]. Even though GSK-3β has been postulated as a tumor suppressor in some types of cancer, this kinase can serve as a pro-oncogene in other malignancies such as pancreas, colorectal, hepatocellular carcinoma, kidney, leukemia, and GBM (reviewed in [28]). Previously, we reported that T98G cells derived from a human GBM contain a functional clock that regulates metabolic oscillations and temporal susceptibility to Bortezomib administration (proteasome inhibitor) [29]. In vivo studies in our laboratory demonstrated higher efficacy in impairing tumor growth when Bortezomib was applied at night in tumor-bearing mice compared to diurnal administration, even at low doses of the drug [30]. Moreover, T98G cultures showed significant differences in cell viability across time when cells were treated with SR9009 (synthetic REV-ERB agonists), an effect that was further potentiated when SR9009 was combined with Bortezomib [31]. Environmental alterations, genetic mutations, or changes in circadian gene expression can disrupt the normal functioning of the circadian machinery and increase the risk of cancer occurrence and progression (reviewed in [32, 33]). In fact, the International Agency for Research on Cancer (IARC) classified “shift work leading to a circadian disruption” as a probable human carcinogen (Group 2A) [34]. For example, studies have shown that the negative element PER2 plays a critical role in GBM growth and metabolism [35]. Furthermore, Ma and colleagues revealed that PER proteins are significantly downregulated in glioma tissues with respect to normal brain samples [36] (reviewed in [24]). In brief, we hypothesize that both the pharmacological modulation of the circadian machinery and the genetic disruption of the molecular clock could severely affect tumor biology, with a strong impact on cell viability. To test this hypothesis and with a view to finding novel strategies to treat GBM, we evaluated the effects of 1) synthetic inhibitors of cytosolic kinases such as CHIR99021 (GSK-3 inhibitor) and PF670462 (CK1ε/δ inhibitor), 2) the cryptochrome protein stabilizer (KL001), or 3) clock gene *Per*2 disruption, on cell viability, metabolic state, cell cycle distribution, and migration in T98G cells.

## RESULTS

### Circadian clock modulation and genetic disruption implications for GBM cell viability and migration

#### Pharmacological modulation: PF670462, CHIR99021, and KL001 treatment

To investigate whether pharmacological modulation of the circadian clock affects GBM viability, T98G cells were treated with a CK1ε/δ inhibitor (PF670462), a GSK-3 inhibitor (CHIR99021), or a CRY protein stabilizer (KL001) in a range of concentrations from 0.01 to 100 μM for 48 hours. The dose-response curve evidenced a half-maximal inhibitory concentration (IC50) of 1.5 μM (R^2^=0.96), 8.6 μM (R^2^=0.94), and 25.4 μM (R^2^=0.96) for PF670462, CHIR99021, and KL001, respectively (Fig. 1). The value of R^2^ denotes the goodness of fit of the experimental data with a symmetrical sigmoidal curve. The determined IC50 values were used to perform subsequent experiments. To elucidate whether pharmacological modulation impairs tumor proliferation and migration, wound-healing experiments were performed after treatment of T98G cells with the CK1ε/δ inhibitor, the GSK-3 inhibitor, or the CRY protein stabilizer at the mentioned IC50 for 24 h. Results showed that vehicle-treated cells had covered about 65.1 % of the wound 24 h after the scratch whereas cells incubated with the inhibitor CHIR99021 filled only 35% of the gap (p < 0.021 by unpaired t-test). On the contrary, no significant differences were observed in the percentage of wound closure between T98G cells treated with the CK1ε/δ inhibitor or the CRY protein stabilizer and control cultures (Fig 2 a-b). Interestingly, lower expression levels of vimentin, an intermediate filament protein implicated in migration and epithelial-mesenchymal transition of tumor cells, were observed in CHIR99021-treated cells (p < 0.0088 by unpaired t-test) (Suppl. Fig. 2 a-b). In another series of experiments, we evaluated the effect of SR9009 treatment on T98G migration. By acting as a REV-ERBs-specific agonist, this synthetic pyrrole derivative is able to modulate the circadian clock function. Previously, our laboratory demonstrated that SR9009 treatment decreased T98G cell viability and impaired cell cycle progression [31]. In this case, the percentage of wound closure of SR9009-treated cells (∼26%) was significantly lower than that observed in cultures incubated with the vehicle (∼57%) (p < 0,006 by Mann Whitney test) (Fig. 2 c-d).

**Fig. 1:**
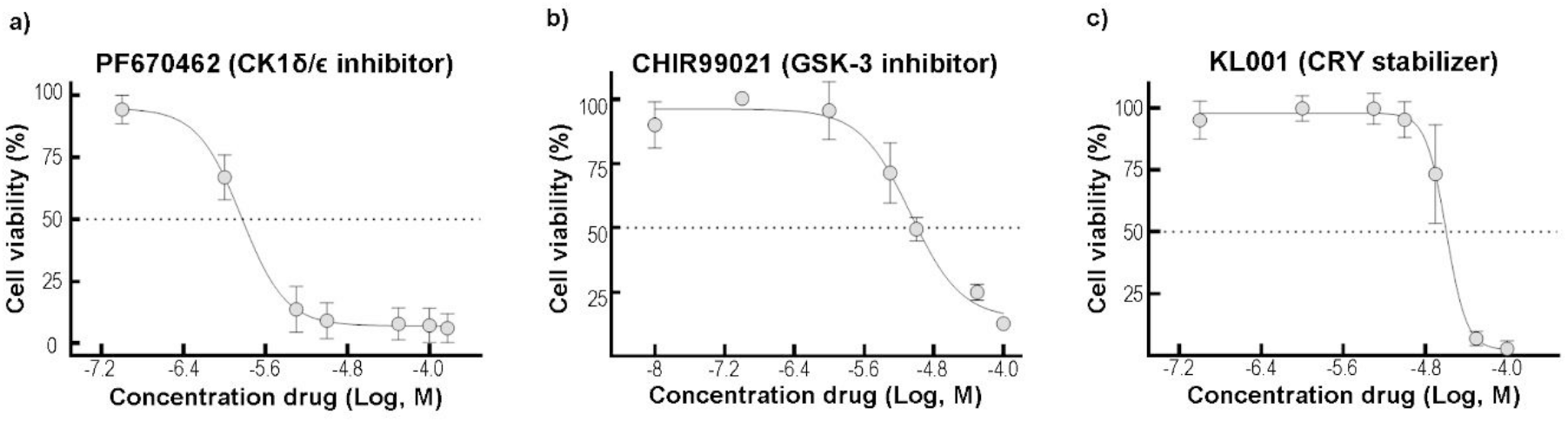
Effect of circadian pharmacological treatment on GBM cell viability. Dose-response curve of PF670462 **(a)**, CHIR99021 **(b)**, and KL001 **(c)** in T98G cells. Culture cells were treated with a range of concentrations (0.01, 0.1, 1, 5, 10, 50, and 100 μM) of the pharmacological modulators for 48 h. Cell viability was measured by alamarBlue assay and the results analyzed by GraphPad Prisma software showed an IC50 value of 1.5 μM (R=0.96), 8.6 μM (R=0.945), and 25.4 μM (R=0.96) for PF670462, CHIR99021, and KL001, respectively. The results are mean ± SD of two independent experiments (n=3/group).

**Fig. 2:**
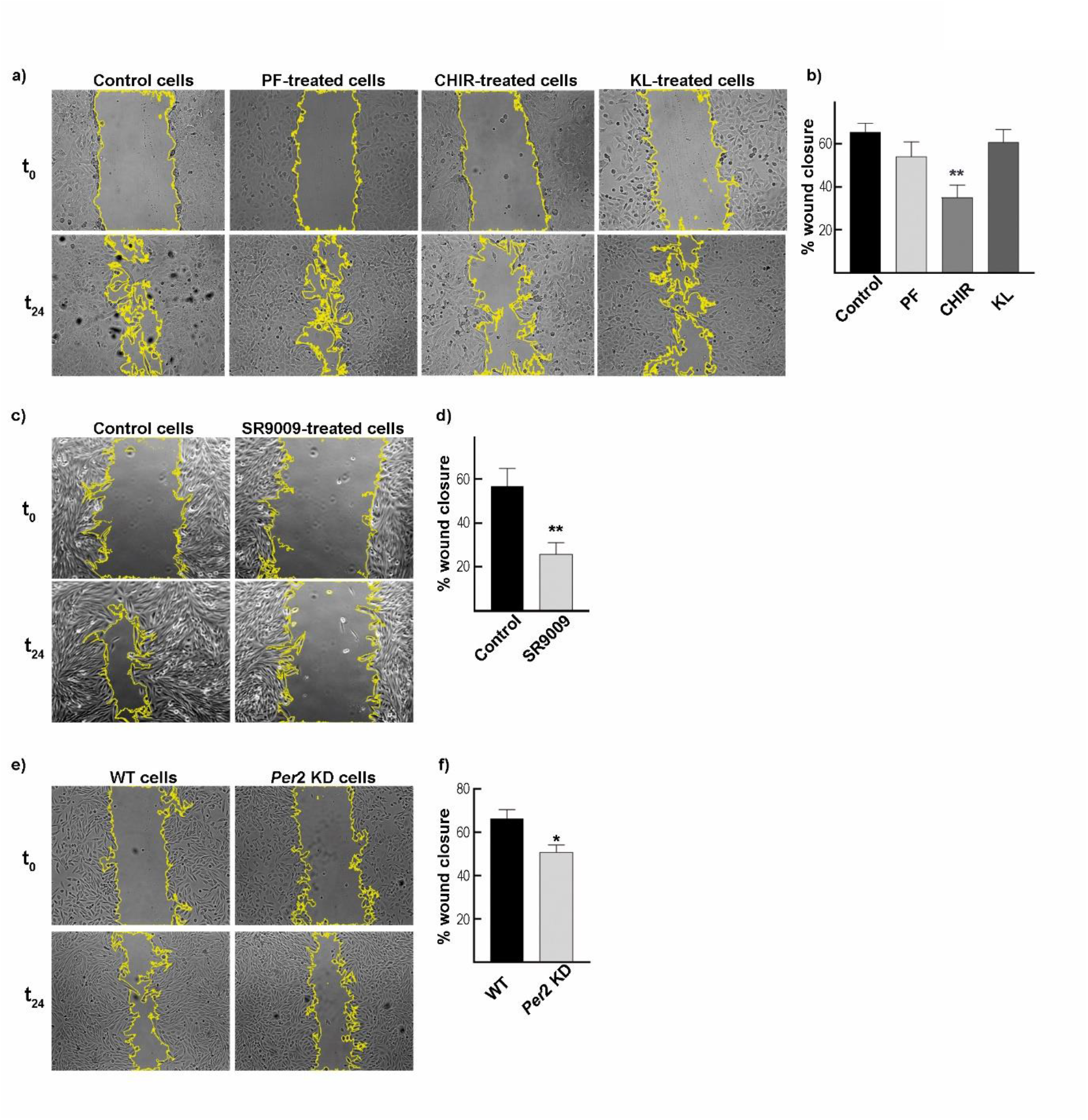
Effect of the pharmacological modulation and *Per*2 knockdown (KD) on GBM cell proliferation and migration. **a)** Representative microphotographs of the scratch done on the monolayer of T98G incubated with PF670462 (1.5 μM), CHIR99021 (8.6 μM), or KL001 (25.4 μM) were taken at t_0_ and t_24_. Control cells were incubated with DMSO (vehicle). **b)** Wound closure percentage showed a significant decrease in the covered area of T98G incubated with CHIR99021 as compared to vehicle-treated cells (p < 0.021 by unpaired t-test) 24 h after the scratch. No significant differences were observed in the percentage of wound closure between PF670462 and KL001 compared to vehicle-treated cells. See Materials and Methods section for further details. PF: PF67046; CHIR: CHIR99021; KL: KL001. **c)** Representative microphotographs of the scratch on SR9009-treated (20 μM) and control cells were taken at t_0_ and t_24_. Control cells were incubated with DMSO (vehicle). **d)** Wound closure percentage showed a significant decrease in the covered area of SR9009-treated T98G cells compared to vehicle-treated cells 24 h after the scratch (p < 0.006 by Man Whitney test). **e)** Representative microphotographs of the scratch done on the monolayer of *Per*2 KD and WT T98G cells were taken at t_0_ and t_24_. **f)** Wound closure percentage showed a significant decrease in the covered area of *Per*2 KD T98G cells as compared to WT cultures (p < 0.021 by Man Whitney test) 24 h after scratch. The results are mean ± SEM of two/three independent experiments performed in triplicate. *p < 0.05, **p < 0.01.

#### Genetic disruption: Per2 knockdown model

In order to determine whether disruption of the molecular clock affects GBM cell proliferation and migration, wound healing experiments were carried out in WT cells or *Per*2 knockdown (KD) cultures. The results indicated that cells with altered expression of PER2 filled only 51% of the gap compared with 66% of wound closure observed in WT cultures (p < 0,021 by Mann Whitney test) (Fig. 2 e-f).

### Cell cycle distribution after CHIR99021 treatment or *Per*2 disruption

To explore whether the results of cell viability and migration could be linked to cell cycle distribution, T98G cells were treated with the GSK-3 inhibitor for 24 h, fixed, and stained with propidium iodide. Flow cytometry results showed a higher percentage of cells in the sub-G_0_ phase on CHIR99021-treated (∼6%) cells compared to control cultures (∼1%) (p < 0.01 by unpaired t-test). The percentage of cells in the G_0_-G_1_ phase among treated cells was significantly lower (∼29%) than in vehicle-treated cells (∼51%) (p < 0,0001 by unpaired t-test), whereas the S-phase showed a significant higher proportion of cells in CHIR99021-treated cultures (∼53%) than in control cells (∼38%) (p < 0,0014 by unpaired t-test). No significant differences were observed in the G_2_-M phase among the experimental conditions (Fig. 3 a-b). Since *Period* genes have been shown to be downregulated in glioma samples compared to normal brains [36, 37], we evaluated how the disruption of *Per*2 affects cell cycle progression. For this, WT and *Per*2 KD cells were arrested in FBS-free DMEM for 48 h and then stimulated with 20% FBS in DMEM medium allowing cultures to re-enter to cell cycle. Significant differences were observed in the G_0_-G_1_ phase with 52 % and 27 % of cells in WT and *Per*2-KD cultures, respectively (p < 0,0002 by unpaired t-test). By contrast, a higher percentage of cells were found in the S-phase (49%) after *Per*2 disruption than in the WT condition (29%) (p < 0,003 by unpaired t-test). However, no significant differences were found in the sub-G_0_ or G_2_/M phases between the experimental conditions examined (Fig 3 c-d). The results obtained may reflect a greater proliferative proportion of *Per2* KD cells, likely due to cells retained in the S-phase.

**Fig. 3:**
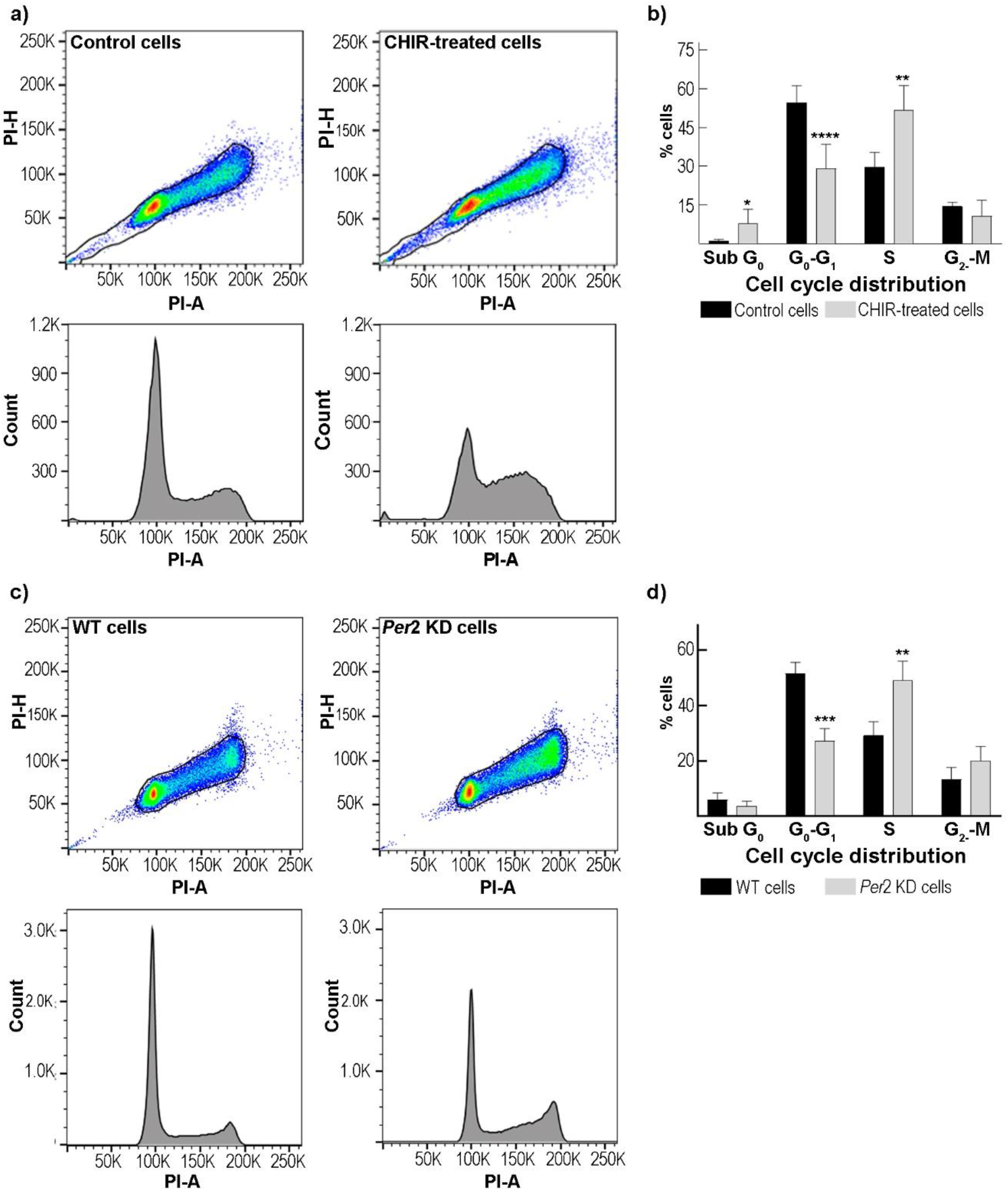
GSK3 inhibitor treatment and *Per*2 downregulation effects on GBM cell cycle progression. **a)** Representative dot plots and histograms of cell cycle distribution using FlowJo. T98G cells were incubated with CHIR99021 (8.6 μM) or vehicle for 24 h and then fixed with ethanol 70% and stained with propidium iodide. **b)** Histograms of the percentage of cells in each phase showed a higher proportion of CHIR99021-treated cells in the sub-G_0_ and S phase and a decrease in the G_0_-G_1_ phase compared to control cultures. No significant differences were observed in the proportion of cells in the G_2_-M phase between the experimental conditions. The results are mean ± SEM of three independent experiments performed in triplicate. *p<.0.05, **p<0.01, and ***p<.0.001 by unpaired t-test. **c)** Representative cell cycle distributions of WT and *Per*2 KD T98G cells. Cultures were arrested in serum-free DMEM for 48 h and then stimulated with 20% FBS for 16 h. Cell cycle phases were analyzed by staining with propidium iodide and flow cytometry. **d)** Quantification of cell cycle distribution showed a lower percentage of *Per*2 KD T98G cells in the G_0_-G_1_ phase and a higher proportion in the S phase with respect to WT cultures. **p<0.01, ***p<0.001 by unpaired t-test. The results are mean ± SEM of three independent experiments performed in triplicate.

### Characterization of clock gene expression on glioblastoma cells after CHIR99021 treatment or *Per*2 disruption

To further investigate the effect of pharmacological modulation on the circadian machinery, T98G cells were treated with the IC50 of CHIR99021 for 24 h, and clock protein expression was evaluated by immunocytochemistry. Results evidenced higher protein levels of the clock activator BMAL1 in cells treated with CHIR99021 compared to control cultures both in the nucleus (p < 0.028 by Mann-Whitney test) and in the cytoplasm (p < 0.0244 by unpaired t-test). On the contrary, levels of REV-ERBα protein showed a significantly higher expression only in the nucleus of CHIR99021-treated cells as compared with vehicle-incubated cells (p < 0.0001 by unpaired t-test) (Fig 4 a-b). Similar results were observed by western blot (Suppl. Fig. 2 a-b), which showed higher levels of BMAL1 protein in T98G-treated cells with the GSK-3 inhibitor. On the other hand, higher levels of PER1 protein were observed by immunocytochemistry in T98G cells after *Per*2 disruption with respect to WT cells (p < 0.0011 by unpaired t-test) (Fig 4 c-d), likely indicating a compensatory effect on *Per1* or the lack of the repressor activity after *Per2* KD. Additional effects were found on the expression of the clock activator *Bmal*1 by PCR (Suppl. Fig. 1 b).

**Fig. 4:**
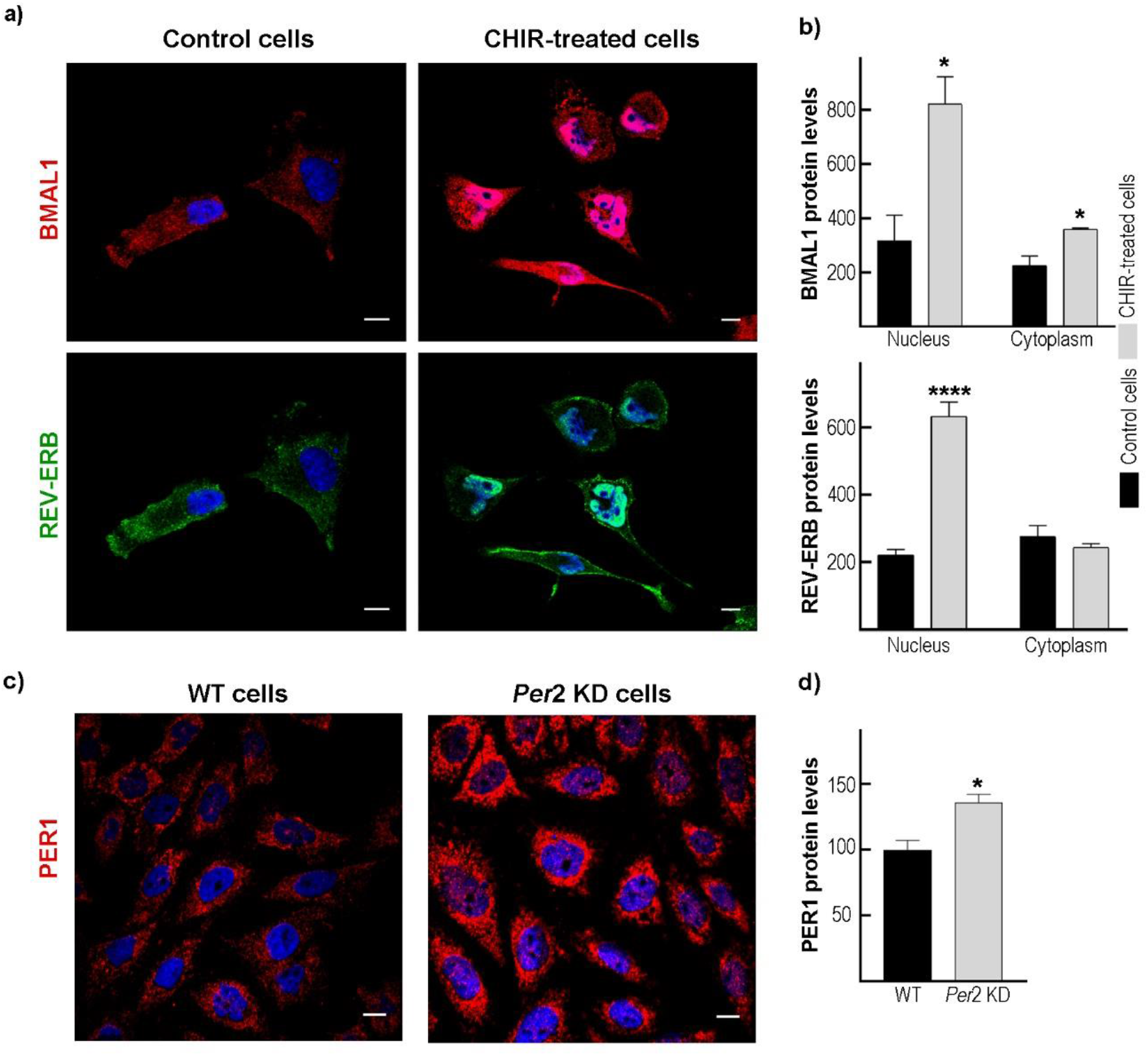
Characterization of clock genes expression after CHIR treatment and *Per*2 knockdown on GBM cells. **a)** Representative microphotographs of CHIR99021-treated cells (right column) for 24 h or control cultures (left column) immunolabeled with antibodies for BMAL1 (red), REV-ERBα (green), and DAPI for nuclear localization. Scale bar = 10 μm. **b)** Quantification of protein levels of BMAL1 (upper graph) showed a higher expression of the clock activator both in the nucleus and cytoplasm in T98G cells treated with CHIR99021 (gray bars). In contrast, REV-ERBα expression (bottom graph) showed higher levels only in the nucleus as compared with control cells (black bars). See Materials and Methods section for further details. *p < 0.05; ***p < 0.0001 by unpaired t-test. **c)** Representative microphotographs of WT (left column) and *Per*2 KD (right column) T98G cells immunolabeled with the antibody for PER1 (red), and DAPI for nuclear localization. Scale bar = 10 μm. **d)** Quantification of PER1 protein levels showed higher expression in *Per*2 KD (gray bar) T98G cells compared to WT cultures (black bar). *p < 0.05 by unpaired t-test.

### Metabolic alterations after CHIR99021 treatment

#### Redox state

To investigate the ability of the GSK-3 inhibitor to alter the redox state, T98G cells were incubated with CHIR99021 at a final concentration of 8.6 μM for 24 h and reactive oxygen species (ROS) levels were measured using the fluorescent probe 2′,7′-dichlorodihydrofluorescein diacetate. The results analyzed by flow cytometry showed a significant increase in ROS levels in CHIR99021-treated cells with respect to vehicle-treated cells (p < 0.014 by unpaired t-test). Interestingly, a higher percentage of positive propidium iodide cells was observed in dot plots of flow cytometry in cells incubated with the GSK-3 inhibitor as compared to control cultures (p < 0.004 by Mann Whitney test), strongly indicating positive propidium iodide staining of dead cells (Fig. 5 a-c).

**Fig. 5:**
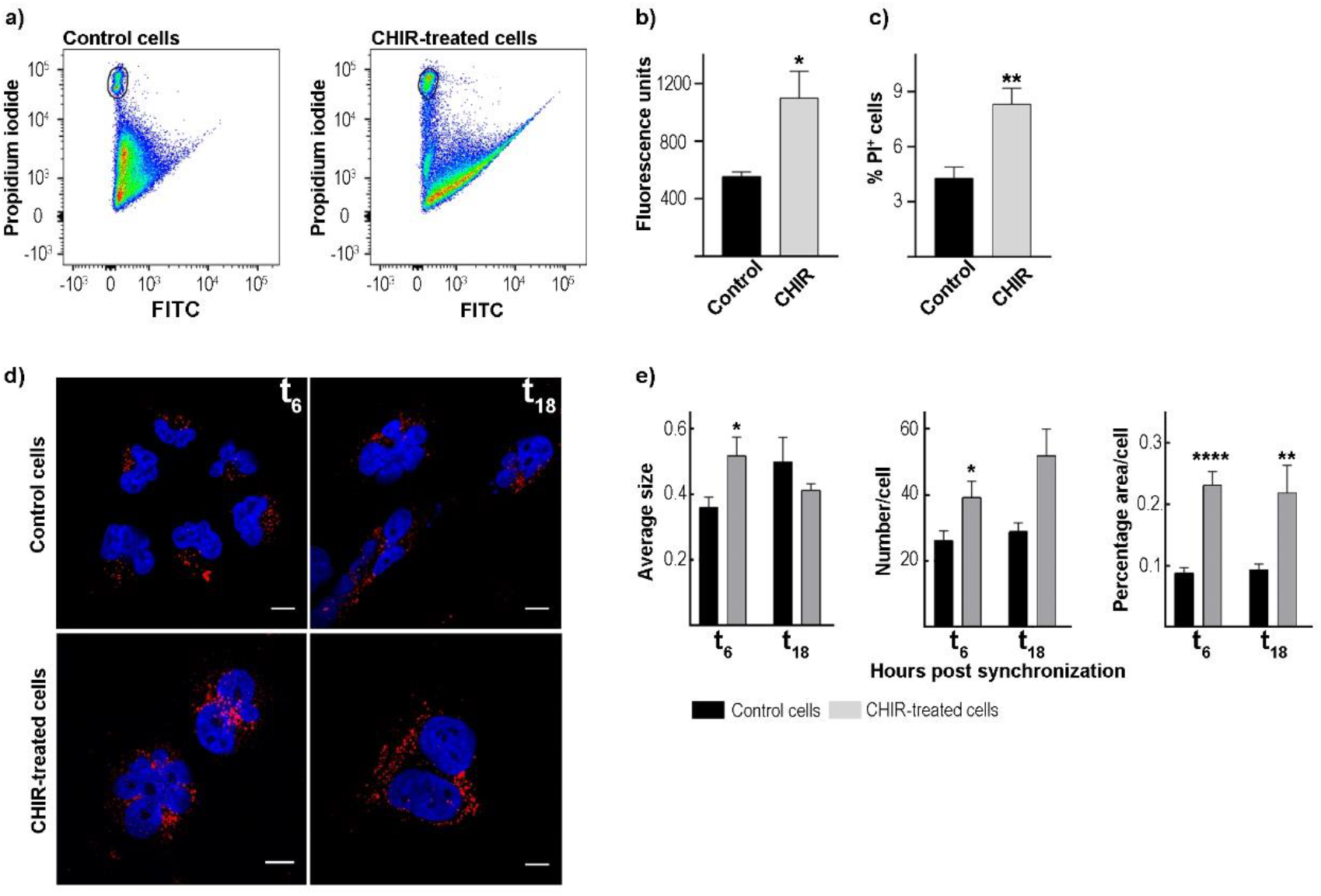
Redox state and lipid droplet determination in GBM cells after GSK-3 inhibition. **a)** Representative dot plots of reactive oxygen species (ROS) levels analyzed in CHIR99021 (8.6 μM) or vehicle-treated cells by flow cytometry with the fluorescent probe 2′,7′-dichlorodihydrofluorescein diacetate. Quantification of fluorescent levels (left column graph, **b**) and positive propidium iodide cells (right column graph, **c**) showed higher levels in cells treated with the GSK-3 inhibitor (gray bars) compared to the control cultures (black bars). The results are mean ± SEM of three independent experiments performed in triplicate. *p < 0.05 by unpaired t-test, **p < 0.01 by Mann Whitney test. PI: propidium iodide, CHIR: CHIR99021. **d)** Representative microphotographs of LD staining in T98G cultures or CHIR99022-treated cells synchronized by a 1-h pulse of DEX (100 mM) and fixed at 6 (left column) and 18 (right column) hours after synchronization. LD fluorescence staining of Bodipy is shown in red, and nuclei stained with DAPI in blue. Scale bar = 10 μm. **d)** Quantification of LD parameters showed a significant increase in average size, percentage area, and number of LDs after GSK-3 inhibition (gray bar) when the cells were fixed at 6 h after synchronization as compared with control cultures (black bar); CHIR99021-treated cells only showed higher levels in terms of LD percentage area with respect to non-treated cells fixed at 18 h after DEX-pulse. *p<.0.05, **p<.0.001, ***p<.0.0001 by unpaired t-test or Mann-Whitney test. CHIR: CHIR99021.

#### Lipid droplets

Considering that BMAL1 and REV-ERB are targets of post-translational modifications by the GSK-3, and these clock proteins are closely implicated in cellular metabolism, we evaluated whether the pharmacological modulation of the circadian machinery could alter the cytosolic/metabolic oscillator. We therefore analyzed different parameters of lipid droplets (LDs), the cytoplasmic organelles mainly involved in energy storage [38–40]. To this end, fixed cells were stained with Bodipy dye, which has highly lipophilic neutral characteristics that allow it to pass through the cell membrane to localize polar lipids and specifically stain LDs [41]. Confocal microscopy results showed a larger average size and number of LDs in T98G cells fixed 6 h after DEX synchronization in CHIR99021-treated cells (0.52 μm^2^ and ∼39 LDs/cell, respectively) as compared to control cultures (0.36 μm^2^ and ∼26 LDs/cell, respectively) (p < 0.02 and p < 0.03 by unpaired t-test, respectively); no significant differences between the two experimental groups were observed at 18 h after synchronization. However, there was a significantly higher percentage area of LDs in T98G cells previously incubated with the GSK-3 inhibitor both at 6 and 18 h after DEX synchronization with respect to control cells (p < 0,0001 by unpaired t-test and p < 0,003 by Mann Whitney, respectively) (Fig. 5 d-e). These results strongly indicate a significant effect of the GSK-3 inhibitor on LD levels at the times evaluated.

### Temporal susceptibility to combined treatment with temozolomide and CHIR99021

Given the effects of CHIR99021 treatment in T98G cells in terms of cell viability and metabolic state, we further investigated whether this pharmacological modulation of the circadian clock could serve to improve the standard temozolomide (TMZ) treatment for GBM patients. To this end, T98G cells were synchronized and treated with CHIR99021 (8.6 μM), TMZ (844 μM), or a combination of both drugs at 6 or 18 h after synchronization for 48 h. The IC50 of TMZ was previously determined in our laboratory (data not shown). The MTT assay results show a significant reduction in cell viability (∼44%) with the combined treatment at 18 h after DEX synchronization compared with the effect of each drug alone (p < 0.05 for TMZ vs TMZ+CHIR and p < 0.01 for CHIR vs TMZ+CHIR by one-way ANOVA) (Fig 6 b). By contrast, the combined treatment applied at 6 h after synchronization only showed a significant decrease in cell viability with respect to those cells incubated with CHIR99021 treatment alone (p < 0.01 by one-way ANOVA) (Fig 6 a). These results indicate a significant temporal variation in combined chemotherapy treatment.

**Fig. 6:**
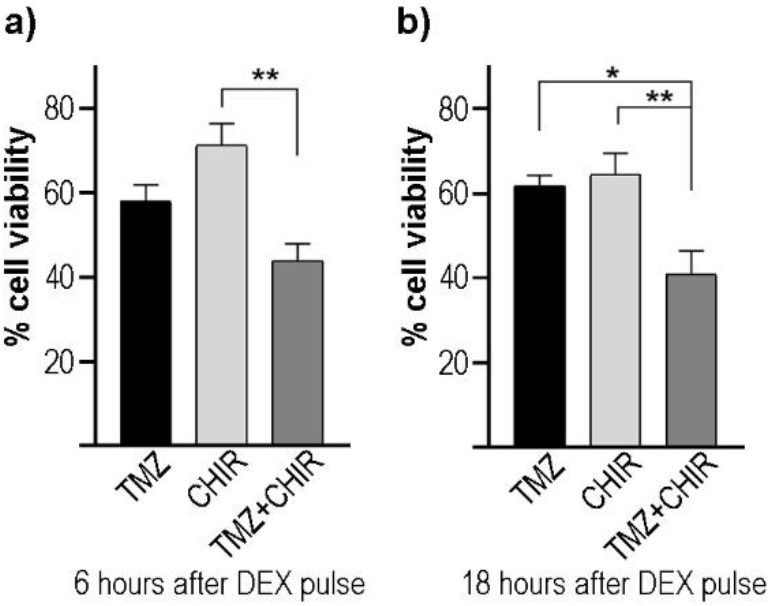
Temporal cell susceptibility to combined treatment with TMZ and CHIR99021. T98G cells were synchronized by a 1-h pulse of DEX (100 nM) and treated with the drugs at 6 and 18 h after synchronization for 48 h. Cell viability analyzed by MTT assay revealed a significant effect when cells were treated with TMZ (844 μM) and CHIR99021 (8.6 μM) at 18 h after synchronization compared to treatment with each drug alone **(b)**. On the contrary, the combination of drugs at 6 h after DEX synchronization showed a significant reduction in cell viability only compared to CHIR99021-treated cells **(a)**. The results are mean ± SEM of four independent experiments performed in triplicate. *p < 0.05, **p < 0.01 by one-way ANOVA.

## DISCUSSION

GBM is a highly vascularized and invasive brain tumor with a poor prognosis. Median patient survival after diagnosis is 12-15 months and there are no full curative therapies (reviewed in [42]). Recent evidence in the field shows that circadian clock function should be considered a novel target to design new therapeutic strategies or improve the current approaches to treat high-grade gliomas. We thus investigated the effects of pharmacological modulation of the circadian clock in terms of cell viability, proliferation, and migration on GBM cultures using the CK1ε/δ inhibitor (PF670462), the GSK-3 inhibitor (CHIR99021), and the CRY protein stabilizer (KL001). Next, we selected the CHIR99021 inhibitor to evaluate cell cycle distribution, clock protein expression, metabolic parameters, and combined treatment with TMZ. In parallel, we examined the effects of circadian disruption on migration and cell cycle progression in a *Per*2 knockdown GBM model. Dose-response curves using the selective inhibitors of CK1 ε/δ (PF670462), GSK-3 (CHIR99021), and CRY degradation (KL001) showed IC50 in the low micromolar range after 48 h of treatment. Although different GSK-3 inhibitors have been evaluated in GBM models [27, 43–48], our results demonstrate for the first time that CHIR99021 treatment impairs GBM cell viability. The migratory capacity of tumor cells is a characteristic of more aggressive or malignant cancer types. To test this capacity we performed wound healing experiments, which are well-established techniques to determine collective migration that regulates wound repair, cancer invasion, metastasis, immune response, and angiogenesis [49–51]. The results of our wound-healing assay revealed that CHIR99021-treated cells (8.6 μM) filled only 35% of the gap after 24 h of treatment compared to 65.1% of wound closure in control cultures. Lower levels of wound closure were also observed when T98G cells were treated with the synthetic agonists of the nuclear receptors REV-ERB. By contrast, no significant differences were observed in the percentage of wound closure between cultures treated with PF670462 (1.5 μM) or KL001 (25.4 μM) and control cells. In this connection, Lin and collaborators showed that KL001 (12 μM) or PF670462 (1.5 μM) treatment for 24 h did not significantly affect cancer cell migration compared to osteosarcoma non-treated cells [52]. Reports in the literature support our results, suggesting that GSK-3 has a key role in regulating epithelial-mesenchymal transition (EMT) in prostate and bladder tumors (reviewed in [28]). Moreover, the ATP-competitive GSK-3 inhibitor 9-ING-41 impairs the tumorigenicity of renal cancer cells by changing the mesenchymal state to the epithelial direction [53]. EMT induces numerous biochemical changes in epithelial cells to switch to a mesenchymal phenotype, resulting in an enhanced invasive capacity [54]. In GBM, WNT and β-catenin, two components that contribute to mesenchymal transition, are highly expressed and correlated with significantly short survival in GBM patients [55]. Evidence from the laboratory of Richardson and colleagues showed that transcriptional targets of GSK-3β regulate GBM invasion [56]; Nowicki *et al.* further showed that the effect on migration after GSK-3β inhibition occurred at earlier time points than the effects on cell viability [57]. The type III intermediate filament, vimentin, is a key biomarker of EMT which is normally expressed in mesenchymal cells but is upregulated during cancer metastasis (reviewed in [58]). In agreement with this, we reported lower levels of vimentin by western blot after CHIR99021 treatment of T98G cells, highlighting the effect of GSK-3 inhibition in tumor migration.

Since a higher expression and activity of GSK-3β in GBM cultures with respect to normal brain samples [27] has been reported and we described significant effects on cell viability and migration in CHIR99021-treated cells, we decided to examine the cell cycle distribution and metabolic alterations after GSK-3 inhibition in more detail. Uncontrolled proliferation is a hallmark of cancer and GSK-3 has an important role in cell cycle regulation since cyclins are phosphorylated by this kinase [59, 60]. We therefore treated T98G cells with CHIR9009 and stained them with propidium iodide followed by flow cytometry analysis. Our results showed a significant alteration in cell cycle distribution, with an increase in the percentage of cells in the S-phase after GSK-3 inhibition compared to control cells. Histograms showing DNA content also revealed ∼6% and ∼1% of cells in the sub G_0_ phase in the treated condition and control cultures, respectively. These results suggest a higher percentage of dead cells after CHIR99021 treatment, with the increase in the S-phase possibly indicating cell cycle arrest at this phase. In line with this, various studies have reported alterations in cell cycle distribution after GSK-3 inhibition in GBM cell lines and patient-derived GBM cultures [43, 44]. With regard to circadian disruption, propidium iodide-stained cells revealed accumulation in the S-phase in the *Per*2 KD condition after serum stimulation with respect to control cultures. However, no differences were observed between the experimental conditions examined in the count of positive propidium iodide cells. Nevertheless, *Per*2 KD cells also display a significant decrease in wound healing closure as compared to WT cells, strongly indicating that disruption of this clock gene alters the migration, and ultimately the metastatic capacity of these glioma cells.

Given that phosphorylation of most clock proteins occurs rhythmically, alteration of clock post-translational modifications can also alter the circadian periods [61–63]. In particular, BMAL1 phosphorylation by GSK-3 leads to ubiquitination and degradation by proteosoma [64] whereas this post-translational modification in REV-ERBα protein protects it from degradation [65, 66]. Phosphorylation of REV-ERB protein promotes nuclear translocation and inhibition of ROR-mediated *Bmal*1 transcription [67]. In line with this, we observed higher levels of BMAL1 protein by western blot and immunocytochemistry techniques after CHIR99021 treatment. By contrast, GSK-3 inhibition showed higher levels of the nuclear receptor REV-ERB in the nucleus compared to the control condition. Considering that BMAL1 and REV-ERB proteins are critical regulators of cellular metabolism [68–72] (reviewed in [73]), we evaluated the redox state and lipid droplet (LD) content in CHIR99021-treated cells. Flow cytometry showed higher levels of ROS in cultures after treatment with the inhibitor as compared to control cells. ROS are considered byproducts of normal cellular oxidative processes and have been implicated in the initiation of apoptotic signaling [74]. After GSK-3 inhibition our results also clearly show metabolic alteration in GBM cells in terms of LDs. These dynamic organelles are surrounded by a monolayer of polar and amphipathic phospholipids with structural proteins such as the perilipin family [75] and function mainly as regulators of storage and hydrolysis of neutral lipids, which are located in the core of the organelles [75, 76]. LD biology has attracted growing interest in recent years since these organelles are implicated in a plethora of functions such as lipid metabolism, control of protein homeostasis, sequestration of toxic lipid metabolic intermediates, protection from stresses, proliferation of tumors, and platforms for pathogen multiplication and defense (reviewed in [77]). Our findings with Bodipy staining and confocal microscopy reveal a larger average size, number, and percentage area of LDs in CHIR99021-treated cultures with respect to control cells at specific times after synchronization, probably highlighting regulation by the circadian clock. Our findings may reflect either a protective mechanism to detoxify the increase in ROS levels or a metabolic reprogramming mechanism that requires further investigation.

Although TMZ treatment is the standard therapy approved by the FDA, the median survival of GBM patients after diagnosis is 12-15 months and less than 5% of patients survive 5 years [78]. Mouse models and clinical studies show that different GSK-3 inhibitors improved the monotherapy of TMZ [47, 48], probably by silencing the O^6^ -methylguanine DNA methyltransferase [79] which is considered an independent factor in GBM patients [80]. In this report, we show that the combination of CHIR99021 and TMZ improved the effects on cell viability compared to treatment with each drug separately. Our findings underline the importance of designing therapeutic strategies based on the circadian clockwork since the combined treatment of these drugs was more effective when they were applied 18 h after synchronization. In support of these findings, we previously demonstrated temporal susceptibility to treatment with the proteasome inhibitor Bortezomib in both in vitro and in vivo models with a temporal window of higher efficacy between 12 and 24 h after culture synchronization or when chemotherapy was applied at the night phase, respectively [29, 30]. Differential responses to TMZ treatment were also reported by Slat *et al.*, suggesting a *Bmal*1-dependent mechanism [81]. Differences in survival or tumor weight were also observed in tumor-bearing mice in accordance with the drug administration time [82, 83].

Overall, the inhibition of GSK-3 using the ATP-competitive CHIR99021 inhibitor affects GBM viability, migration, and cell cycle distribution as well as clock protein levels and metabolic parameters. Genetic disruption of the clock gene *Per*2 also alters the circadian clock function in migration and cell cycle progression. Although further studies are needed to elucidate the mechanisms involved, our approach could offer a potential therapeutic strategy by using CHIR99021 in combination with TMZ to treat GBM. Based on the literature and our current findings, such a combination will lead to a better understanding of the circadian clock function in health and disease and the design of strategies aimed at maximizing therapy effectiveness with lower side effects.

## CONCLUSIONS

Our findings strongly support the idea that both the pharmacological modulation and genetic disruption of the circadian clock severely alter glioblastoma cell biology. A significant temporal variation in drug susceptibility was observed in the combined chemotherapy treatment, highlighting the role of the circadian clock at specific times in maximizing therapeutic benefits while minimizing side effects. Since metabolic reprogramming is a hallmark of cancer cells, the metabolic alterations reported in this manuscript in combination with the circadian modulation could result in a novel therapeutic approach to improve GBM treatment.

## MATERIALS AND METHODS

### Cell cultures and synchronization protocol

T98G cells are derived from a human GBM (ATCC, Cat. No. CRl-1690, RRUD: CVCL0556, Manassas, VA, USA) and were tested negative for mycoplasma contamination. Cell cultures were grown in Dulbecco’s modified Eagles medium (DMEM) (Gibco, BRL, Invitrogen, Carlsbad, CA, USA) supplemented with 10% fetal bovine serum (FBS) at 37 °C and 5% CO2 according to [29]. To synchronize cells, cultures were incubated with dexamethasone (DEX, 100 nM) for 1 h at 37°C. Then, cells were washed with phosphate-buffered saline (PBS) and collected at different times post-synchronization according to subsequent experiments.

### *Per*2 disruption by CRISPR/Cas9 genetic edition

The human *Per*2 gene was disrupted in T98G cells using the CRISPR/Cas9 genetic editing tool as previously described [29, 84]. Briefly, we designed single guide RNAs to target the exon 1 of the human *Per*2 gene and subcloned it into the PX459 vector (Addgene) to obtain the PX459-*Per*2 plasmid. The primer sequence corresponding to the single guide RNA was 5′ TCCTCGGCTTGAAACGGCGC 3′. T98G wild-type (WT) cells were transfected with Lipofectamine 2000 (Invitrogen) and selected with puromycin (10 μg/mL) for 4 days. PER2 expression on the isolated clone after serial dilution was evaluated by western blot showing to be at least 50% lower than WT cultures. The complete silencing/knockout of the target gene is not feasible in T98G cells since they have a hyperpentaploid chromosome count (Fig Suppl. 1 a-b).

### Pharmacological treatment and determination of cell viability by alamarBlue assay

T98G cells were plated in 96-well plates at a density of 3*10^3^ and were allowed to attach overnight at 37°C. Cultured cells were incubated with PF670462 (CK1ε/δ inhibitor, Santa Cruz), CHIR99021 (GSK-3 Inhibitor XVI, Santa Cruz), or KL001 (Cryptochrome protein stabilizer, Santa Cruz) at different concentrations (0.01, 0.1, 1, 5, 10, 50, and 100 μM) for 48 h at 37°C. PF670462, CHIR99021, and KL001 stock solutions were resuspended in DMSO to a final concentration of 100 mM, 20 mM, and 10 mM, respectively, according to the manufacturer’s instructions. As control cells, cultures were incubated with the percentage of DMSO corresponding to the maximum concentration tested of drugs. The maximum volume of DMSO in cell culture was always <1%. After incubation, the culture medium was removed and alamarBlue® 10% v/v dissolved in DMEM supplemented with 10% FBS was added to culture cells for 2-3 h. Then, the fluorescence was measured at a wavelength of 590 nm in a BioteK microplate reader [85]. The IC50 value was calculated according to the equation “log(inhibitor) vs response – Variable slope (four parameters) of the GraphPad Prism software. The goodness of fit was evaluated through the R^2^ value.

### Wound healing assay

T98G cells were seeded in a 24-well plate at a density of 1*10^5^ cells and incubated overnight at 37°C to create a monolayer of cells. A P200 pipette tip was used to make a scratch in the middle of the well (t_0_) and the cells were washed with PBS to remove the debris. Fresh culture medium (DMEM + 5% FBS) was added with DMSO (vehicle), PF670462 (1.5 μM), CHIR99021 (8.6 μM), KL001 (25.4 μM), or SR9009 (20 μM) and the cells were incubated for 24 h (t_24_). Similar procedures were used with control cells and *Per*2 knockdown T98G cells without the addition of drugs. Images were taken at t_0_ and t_24_ using the Leica DMI 8 microscope and the wound area was measured using the ImageJ Software. The following formula estimated the percentage of wound closure: % = [1 − (wound area at t_24_/wound area at t_0_)] *100%, according to [86].

### Immunocytochemistry

Culture cells were seeded in coverslips at a density of 1*10^4^ cells and allowed to attach overnight at 37°C. T98G cells were treated with DMSO (vehicle) or CHIR99021 (8.6 μM) for 24 h. Then, cells were fixed in 4% paraformaldehyde for 15 minutes and 10 minutes in methanol according to [31]. Coverslips were washed with PBS and incubated with blocking buffer (PBS supplemented with 0.1% bovine serum albumin, 0.1% Tween 20, and 0.1% glycine) for 1 h at room temperature. Primary antibodies (Table 1) were incubated overnight at 4°C and then, coverslips were washed 3 times with PBS. Finally, goat anti-rabbit immunoglobulin G (IgG; Jackson 549 antibody 1:1000) or goat anti-mouse IgG (Jackson 488 antibody 1:1000) were incubated for 1 h at room temperature. Images were visualized by confocal microscopy (FV1200; Olympus) and analyzed with ImageJ software. Cellular nuclei were visualized by DAPI staining.

**Table 1.** Summary of antibodies used, indicating name, host, catalog number, and dilution for immunochemistry (ICC) and western blot (WB).

| Antibody | Host | Catalog | Dilution |
| --- | --- | --- | --- |
| REV-ERBα | Mouse | Santa Cruz Biotechnology sc-100910 (RS-14).<br>RRID:AB_2154647 | 1:100 (ICC)<br>1:200 (WB) |
| BMAL1 | Rabbit | NOVUS, nb-100-2288. RRID:AB_10000794 | 1:200 (ICC)<br>1:500 (WB) |
| α-TUB | Mouse | Merck Millipore, DM1A | 1:500 (WB) |
| Vimentin | Mouse | Sigma-Aldrich V5255. RID:AB_477625 | 1:500 (WB) |
| PER2 | Rabbit | Abcam ab180655. RRID:AB_2630357 | 1:100 (WB) |
| PER1 | Rabbit | Abcam. ab3443. RRID:AB_303805 | 1:100 (ICC) |

### Western Blot

T98G cells treated with DMSO (vehicle) or CHIR99021 (8.6 μM) for 24 h were harvested in radioimmunoprecipitation assay (RIPA) buffer containing protease inhibitor (Sigma). Proteins were separated by 12% SDS-PAGE according to [29]. Primary antibodies described in Table 1 were incubated overnight at 4°C. Then, membranes were washed with PBS-Tween 0.1% and secondary antibodies were incubated for 1 h at room temperature. Finally, Odyssey IR Imager (LI-COR Biosciences) was used to scan the membranes. The densitometry quantification was carried out with ImageJ software.

### Cell cycle distribution analysis

T99G cells were incubated with DMSO (vehicle) or CHIR99021 (8.6 μM) in DMEM + 5% FBS for 24 h. Then, cells were collected by trypsinization, washed with PBS, and fixed with ethanol 70% at -20°C for at least 24 h. Cells were washed with PBS and pellets were resuspended in PBS containing 50 μg/mL propidium iodide and 10 μg/μl RNAse A as reported in [31]. Cell cycle distribution was analyzed by flow cytometry and subsequent analysis by FlowJo software. For *Per*2 knockdown experiments, control or *Per*2-disrupted cells were arrested in serum-free DMEM for 48 h. Then, the medium was removed and cultures were stimulated with 20 % FSB for 20 h [31]. Cell cycle analysis was performed as described above.

### Reactive oxygen species determination

T98G cells were treated with DMSO (vehicle) or CHIR99021 (8.6 μM) for 24 h at 37°C. Then, cells were collected by trypsinization to evaluate the redox state according to [29]. Briefly, collected cells were washed with PBS and incubated with 2′,7′-dichlorodihydrofluorescein diacetate (Sigma) at 2 μM final concentration for 30 minutes at 37°C protected from light. Cells were washed with PBS and the fluorescence intensity was measured at 530 nm when the sample was excited at 485 nm by flow cytometry. The analysis program used was FlowJo software and live cells were discriminated by propidium iodide stain.

### Lipid droplet determination

1*10^4^ cells were seeded in coverslips and synchronized with 100 nM DEX for 1 h at 37°C. Culture cells were maintained with DMEM supplemented with 5% SFB. After DEX pulse (t_0_), cells were collected at 6 and 18 h after synchronization. For lipid droplet (LD) staining, cells were fixed with 4% paraformaldehyde for 15 minutes and washed twice with PBS according to [87, 88]. Then, coverslips were incubated with Bodipy (Sigma cat#790389, maximum excitation/emission wavelength: 493/503 nm) at a final concentration of 2 μM for 30 minutes protected from light. Coverslips were washed twice with PBS and visualized by confocal microscopy at 60X objective (FV1200, Olympus). Cellular nuclei were visualized by DAPI staining. ImageJ software carried out the LD average size, percentage area per cell, and number per cell quantification. For LD determination on CHIR-treated cells, cultures were incubated with CHIR99021 (8.6 μM) for 24 h prior to synchronization and then analyzed as described above.

### Combined chemotherapeutic treatments

3*10^3^ T98G cells seeded in 96-well plates were synchronized with 100 nM DEX for 1 h at 37°C and maintained with DMEM supplemented with 5% SFB. After DEX pulse (t_0_), cells were treated at 6 or 18 h after synchronization with the combination of temozolomide (TMZ, 844 μM) and CHIR99021 (8.6 μM) or each drug alone. After 48 h, cell viability was analyzed MTT assay as described in [29]. T98G cells incubated with DMSO (vehicle) at 6 or 18 h after synchronization were considered as 100% viability.

## ACKNOWLEDGMENTS

Authors are grateful to Mrs. Gabriela Schanner, Drs. Natalia Baez, Carlos Mas, and Maria Cecilia Sampedro for their excellent technical support and to Dr. César Prucca for experimental advice.

## STATEMENTS AND DECLARATIONS

### FUNDING

This work has been supported by Agencia Nacional de Promoción Científica y Técnica (FONCyT, PICT 2017-631, PICT 2020-0613), Consejo Nacional de Investigaciones Científicas y Tecnológicas de la República Argentina (CONICET) (PIP 2014), Secretaría de Ciencia y Tecnología de la Universidad Nacional de Córdoba (SeCyT-UNC, Consolidar 2018-2022).

### CONFLICT OF INTERESTS

All authors declare that they have no competing interests.

### AUTHOR CONTRIBUTION

PMW, MEG designed research; PMW performed research; MEG contribute new reagents/analytic tools; PMW, MEG analyzed data and wrote. All authors read and approved the final manuscript

---

**Suppl. Fig. 1:**
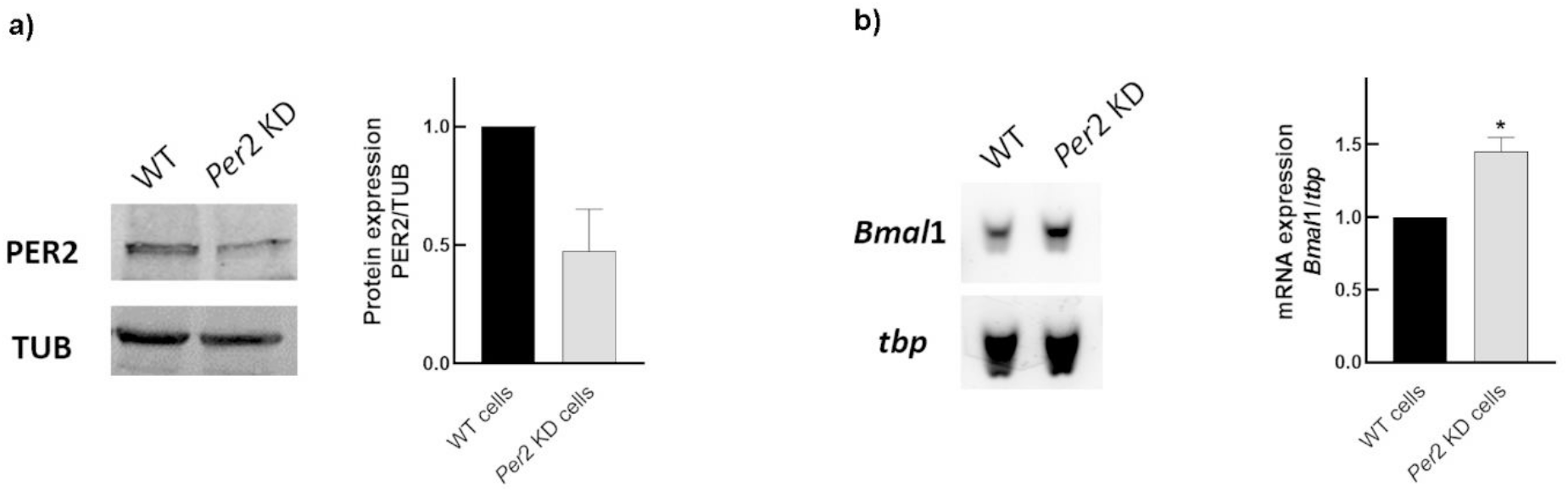
Genetic disruption of the clock gene *Per*2 by CRISPR/Cas9 tool on T98G cells. a) PER2 expression was evaluated by western blot in the isolate clone obtained by serial dilution after transfection and antibiotic selection in T98G cultures. Results showed at least 50% lower levels than non-transfected cultures. Tubulin (TUB) was used as loading control**. b)** *Bmal*1 expression was evaluated by RT-PCR in *Per*2 KD and control cultures. Results showed a higher expression of *Bmal*1 after genetic disruption of *Per*2. Tata binding protein (*tbp)* gene was used as housekeeping.

**Suppl. Fig. 2:**
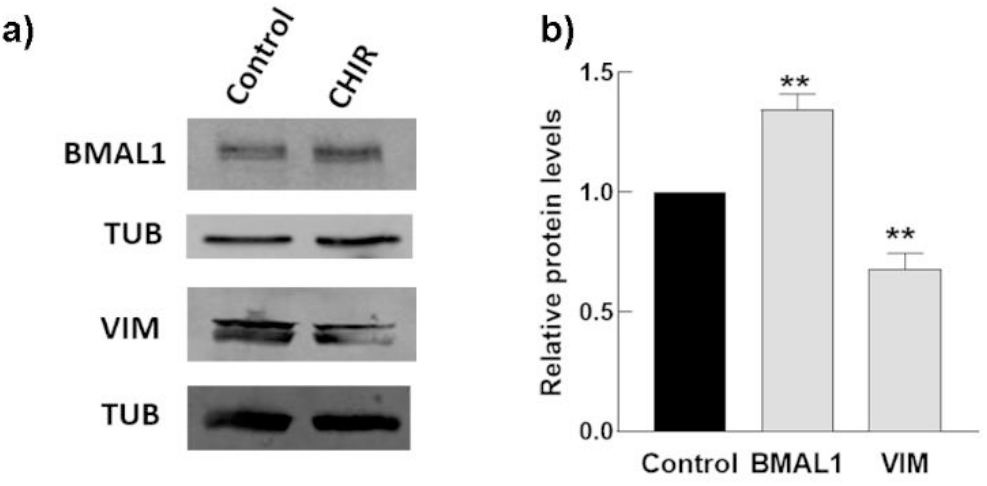
BMAL1 and vimentin expression levels in T98G cells after CHIR99021 treatment. Cultures were incubated with CHIR99021 (8.6 μM) for 24 h and protein levels were detected by western blot. **a)** Representative western blot of BMAL1 and vimentin (VIM) in CHIR99021-treated cells and control cultures. Tubulin (TUB) was used as loading control. **b)** Histograms of relative protein levels indicating an increase in BMAL1 and a decrease in vimentin levels in T98G cells after GSK-3 inhibition (gray bars) as compared to non-treated conditions (black bar). **p < 0.01 by unpaired t-test.

## Notes

### Competing Interest Statement

The authors have declared no competing interest.

